# Prolactin and the shared regulation of parental care and cooperative helping in white-browed sparrow weaver societies

**DOI:** 10.1101/2021.09.22.461403

**Authors:** Lindsay A. Walker, Linda Tschirren, Jennifer E. York, Peter J. Sharp, Simone L. Meddle, Andrew J. Young

## Abstract

In many cooperatively breeding species non-breeding individuals help to rear the offspring of breeders. The physiological mechanisms that regulate such cooperative helping behavior are poorly understood, but may have been co-opted, during the evolution of cooperative breeding, from pre-existing mechanisms that regulated parental care. Key among these may be a role for prolactin. Here we investigate whether natural variation in circulating prolactin levels predicts both parental and helper contributions to nestling provisioning in cooperatively breeding white-browed sparrow weavers, *Plocepasser mahali*. In sparrow weaver groups, a single dominant pair monopolize reproduction and non-breeding subordinates help with nestling feeding. We show that: (i) among parents, dominant females feed nestlings at higher rates, make longer provisioning visits, and have higher prolactin levels than dominant males; and (ii) among subordinates, engaged in cooperative helping behavior, those within their natal groups feed nestlings at higher rates and have higher prolactin levels than immigrants. Accordingly, continuous variation in prolactin levels positively predicts nestling-provisioning rates and mean provisioning visit durations when all bird classes are combined. These relationships are principally driven by differences among bird classes in both circulating prolactin levels and provisioning traits. The more limited within-class variation in prolactin and provisioning traits were not evidently correlated, highlighting a likely role for additional mechanisms in the fine-scale regulation of care. Our findings broadly support the hypothesis that parental care and cooperative helping behavior are regulated by a common underlying mechanism and highlight the need for experimentation to now establish the causality of any role for prolactin.

## INTRODUCTION

In many cooperatively breeding species, non-breeding helpers assist with the rearing of parents’ young, via cooperative contributions to diverse forms of care (e.g. incubation, babysitting and offspring provisioning; Solomon and French, 1997; Koenig and Dickinson, 2004; 2016). The majority of research on such alloparental ‘helping behavior’ has sought to explain its evolution, by identifying the effects of helping on recipients and the means by which these yield fitness benefits to helpers (Cockburn, 1998; Dickinson and Hatchwell, 2004; Koenig and Dickinson, 2016; Capilla-Lasheras et al., 2021). By contrast, our understanding of the proximate physiological mechanisms that regulate the expression of cooperative behavior is less advanced (Schoech et al., 2004; Soares et al., 2010; Sanderson et al., 2014; Dantzer et al., 2017; Vullioud et al., 2021), despite a surge of interest in the origins of consistent individual differences in both cooperative behavior and endocrine traits (English et al., 2010; Sanderson et al., 2015; Dantzer et al., 2019; Houslay et al., 2019; 2022). As cooperatively breeding species typically evolved from species in which parental care was already well developed in both sexes (Cornwallis et al., 2010; Lukas and Clutton-Brock, 2012), it seems likely that the physiological mechanisms that regulate cooperative helping behavior among non-breeders were co-opted from the pre-existing mechanisms that regulated parental care among breeders. Attempts to identify the proximate mechanisms that regulate helping behavior may therefore be well served by testing candidate mechanisms already identified for the regulation of parental care in non-cooperative species (Ziegler, 2000; Schoech et al., 2004; Carlson et al., 2006a; 2006b). One such mechanism is the neuroendocrine pathway involving the anterior pituitary gland hormone prolactin (Buntin, 1996; Sharp et al., 1998; Ziegler, 2000; Carlson et al., 2006a; 2006b; Angelier et al., 2016).

Numerous studies suggest that prolactin can play a causal role in the expression of parental care, though its precise role is not clear and seems likely to vary across taxa (Buntin, 1996; Sharp et al., 1998; Angelier et al., 2016). In birds, prolactin is thought to play a causal role in the onset and maintenance of parental care, but it is less clear whether variation in circulating prolactin levels is also involved in the quantitative regulation of contributions to care once caring behavior has begun (Boos et al., 2007; Angelier et al., 2016; Smiley and Adkins-Regan, 2018). The transition from sexual activity to parenting is typically associated with an increase in circulating prolactin levels, which peak during the care period (Buntin, 1996; Sharp et al., 1998; Angelier et al., 2016; e.g. Schoech et al., 1996; Badyaev and Duckworth, 2005). Evidence that naturally low prolactin levels are commonly associated with breeding attempt abandonment and/or failure (e.g. Chastel and Lormee, 2002; Chastel et al., 2005), and that experimental reductions in circulating prolactin levels can disrupt incubation behavior (e.g. Thierry et al., 2013) and the expression of post-natal nestling care (e.g. Smiley and Adkins-Regan, 2018), suggest that these elevated prolactin levels are necessary for the onset and/or maintenance of both pre- and post-natal parental care. Indeed, experimental elevations of circulating prolactin suggest that elevated prolactin levels can promote the onset of both incubation behavior (e.g. Sockman et al., 2000) and nestling provisioning behavior (e.g. Badyaev and Duckworth, 2005; Buntin et al., 1991). Positive associations between continuous variation in circulating prolactin levels and the rates at which parents provision their offspring (e.g. Duckworth et al., 2003; Ouyang et al., 2011) highlight the possibility that prolactin levels also regulate the *amount* of care that an actively caring parent provides to its offspring. However, such positive associations could arise instead via effects of provisioning activity on a bird’s circulating prolactin levels, as parental contact with offspring cues can increase prolactin secretion (Hall, 1987; Sharp et al., 1998). Causal links between prolactin and provisioning rates could therefore exist in both directions. Indeed, such a feedback loop (in which offspring cues stimulate prolactin secretion that in turn maintains and/or elevates the expression of parental care) could conceivably both maintain parental care while offspring survive, and regulate its expression according to offspring vigor and need (Angelier et al., 2016).

A number of studies of cooperatively breeding species have now begun to investigate the relationships between prolactin and care-giving behavior, both among parents and non-breeding helpers (Ziegler, 2000; Schoech et al., 2004; Soares et al., 2010). Prolactin levels have been shown to rise in parents and non-breeding helpers during the transition to incubation and nestling care in at least three species of cooperatively breeding bird (Schoech et al., 1996; Brown and Vleck, 1998; Khan et al., 2001; see also Vleck et al., 1991). While few studies have investigated specifically whether variation in circulating prolactin levels predicts variation in cooperative contributions to helping, studies of at least two cooperative breeders have yielded compelling evidence in this regard. In Florida scrub jays (*Aphelocoma coerulescens*), breeders fed offspring at higher rates than non-breeders and showed higher circulating prolactin levels (Schoech et al., 1996; see also Vleck et al., 1991), and those non-breeders that helped to feed offspring showed higher prolactin levels than those that did not (Schoech et al., 1996). Continuous variation in circulating prolactin levels also predicted continuous variation in feeding contributions, both among all birds combined and specifically among non-breeders (Schoech et al., 1996). Similarly, in meerkat (*Suricata suricatta*) societies, continuous variation in the prolactin levels of helpers positively predicted their cooperative contributions to both babysitting and pup-feeding (Carlson et al., 2006a; 2006b). In the pup-feeding study, prolactin levels only predicted the cooperative pup-feeding rates of helpers in statistical models that did not allow for an independent positive effect of circulating cortisol levels on the helper’s pup-feeding rates (Carlson et al., 2006a). Experimental work since highlights that this putative positive effect of cortisol on pup-feeding rates may not have been causal, however, as glucocorticoid receptor blockade increased rather than decreased pup-feeding rates among meerkat helpers (Dantzer et al., 2017). Such relationships between circulating prolactin levels and helping behavior are not always apparent. For example, prolactin levels did not predict the offspring provisioning rates of helpers in red-cockaded woodpecker (*Picoides borealis*) groups (Khan et al., 2001), and the pituitary gland prolactin mRNA levels of a cooperatively breeding fish were not evidently related to care-giving behavior (Bender et al., 2008); though the relevant sample sizes in both studies were modest. Attempts to identify correlations between natural continuous variation in prolactin levels and care-giving behavior are expected to be complicated, however, by the existence of other neuroendocrine pathways that may also modulate the expression of care-giving (e.g. glucocorticoids may play a role in its state-dependent modulation; Sanderson et al., 2014; Dantzer et al., 2017; see also Angelier and Chastel, 2009; Angelier et al., 2016), potentially independent of circulating prolactin levels (Schoech et al., 1998; Ziegler, 2000; Young et al., 2005; Carlson et al., 2006a). Nevertheless, the promising findings to date highlight the need for further studies to investigate whether natural circulating levels of prolactin predict variation in contributions to both parental care and helping behavior in cooperatively breeding species, and, ultimately, the use of experimental manipulations to test the causality and nature of any hormone-behavior relationships detected (Sockman et al., 2000; Carlson et al., 2003; Badyaev and Duckworth, 2005; Smiley and Adkins-Regan, 2018).

Here we investigate whether natural variation in circulating prolactin levels positively predicts the nestling provisioning behavior of both parents and non-breeding helpers in a wild cooperatively breeding bird, the white-browed sparrow-weaver (*Plocepasser mahali*). White-browed sparrow weavers are rain-dependent breeders that live in year-round territorial groups throughout the semi-arid regions of sub-Saharan Africa (Lewis, 1982; Wood et al., 2021). Within each social group, a single dominant male and female completely monopolize within-group reproduction and up to 10 non-breeding subordinates of both sexes help to feed their nestlings (Harrison et al., 2013a; 2013b; Capilla-Lasheras et al., 2021; 2023). Subordinates are typically offspring from previous broods that have delayed dispersal from their natal group (and so are helping to rear their parents’ young), but subordinate immigrants of both sexes do also occur (Harrison et al., 2013a; 2013b; Harrison et al., 2014). Subordinates contribute to several cooperative activities year-round, including territorial defense, roost construction and anti-predator vigilance (Lewis, 1982; Walker et al., 2016; York et al., 2019), and during breeding periods they contribute substantially to nestling provisioning (Cram et al., 2015a; Capilla-Lasheras et al., 2021; 2023). Helping behavior by subordinates in this species appears to have positive effects on both the helped offspring being fed and their parents. First, helping behavior has a causal positive effect on the overall rate at which nestlings are fed (Capilla-Lasheras et al., 2021), and, accordingly, helper numbers positively predict nestling survival to fledging during dry periods, reducing the environmentally-induced variance in the reproductive success of the dominant pair (Capilla-Lasheras et al., 2021). Evidence suggestive of positive helper effects on offspring telomere lengths suggest that helpers may also improve the downstream performance of surviving young (Wood, 2017; Wood and Young, 2019). Second, helping behavior appears to lighten the post-natal provisioning workload of the dominant female (Capilla-Lasheras et al., in press), which may explain why mothers with more help increase their pre-natal investment in the egg (Capilla-Lasheras et al., in press) and show higher overwinter survival (O’Callaghan, 2021). While the neuroendocrine correlates of white-browed sparrow weaver reproduction, aggression and song production have been investigated (e.g. Wingfield and Lewis, 1993; Voigt et al., 2007; York, 2012; York et al., 2016), the regulation of parental and helper contributions to offspring provisioning remains unexplored.

We test three predictions of the hypothesis that prolactin promotes the expression of both parental care (among dominants) and cooperative helping behavior (among non-breeding subordinates) in cooperatively breeding societies. First, with regard to parental care, we predict that differences between the nestling provisioning rates of dominant females and dominant males will be mirrored by parallel differences in their mean circulating prolactin levels. Dominant females are expected to provision nestlings at higher rates (and to have higher prolactin levels) than dominant males, as the 12-18% incidence of extra-group paternity in this population leaves dominant females more closely related than dominant males, on average, to the offspring that they rear (Harrison et al., 2013a). Second, with regard to alloparental helping behavior, we predict that differences between the nestling provisioning rates of subordinates still residing within their natal group (hereafter ‘natal subordinates’) and immigrant subordinates, will also be mirrored by parallel differences in their mean circulating prolactin levels. Natal subordinates are expected to provision nestlings at higher rates (and to have higher prolactin levels) than immigrant subordinates, as while the former are typically rearing future generations of siblings born to their parents, the latter are typically unrelated to the nestlings in their group (Harrison et al., 2013a). Finally, we predict that continuous variation in circulating prolactin levels will positively predict continuous variation in the nestling provisioning rates of birds, and that this relationship will be apparent (i) at the population level (when all four of the bird classes above are combined), and (ii) within bird classes, having factored out the among-class differences in prolactin levels and provisioning rates.

## METHODS

### General field methods

Data were collected in the context of a long-term research project that monitors ∼40 cooperative groups of white-browed sparrow weavers at Tswalu Kalahari Reserve, South Africa (27°160S, 22°250E), and at a similar time in two separate breeding seasons (January to February 2013, and January to March 2014). White-browed sparrow weavers in this population may breed at any time from September through to May (the Southern summer), depending on the timing of unpredictable summer rainfall (Wood et al., 2021; Capilla-Lasheras et al., 2021). Each bird within our study population is fitted with a metal ring and three color rings, providing a unique ring combination for identification in the field (SAFRING license 1444). From around six months of age, males and females of the focal subspecies (*Plocepasser mahali mahali*) can be distinguished by their bill color; males have a dark brown bill while females have a paler grey-to-pink bill (Leitner et al., 2009). Dominance status and social group compositions were determined via regular (at least twice weekly) group visits. Social dominance was assigned based on the monitoring of key dominance-related behaviors: the dominant pair routinely displace other group members and produce synchronized duet song, the dominant female is the sole incubator, and the dominant male consistently produces dawn song during breeding periods (Harrison et al., 2013a; Cram et al., 2015b; York et al., 2016). The dispersal status (natal or immigrant) of subordinate birds was determined via the continuous monitoring of the study population since 2007. Based on this information, four classes of birds were assigned: Dominant Females; Dominant Males; Natal Subordinates and Immigrant Subordinates. Social group size was defined as the number of adult (> 1 year of age) birds consistently seen foraging and roosting together at the time of the focal breeding attempt. The breeding status of each group was determined by monitoring the contents of all woven nest structures within each group’s territory, at least every other day throughout the two study periods. When one or more eggs were newly detected, the active nest was visited daily in the afternoon until no new eggs were detected (the birds lay one egg per day in the morning, and typically lay clutches of 2 eggs (range 1-4); Harrison et al., 2013a). To determine hatch dates, daily monitoring of the active nest resumed 14 days after the detection of the first egg (as incubation lasts 14-19 days; Harrison et al., 2013a). This method yielded accurate information on the day on which the first nestling in each clutch hatched, which was termed ‘Day 1’ of the nestling provisioning period for the focal breeding attempt. All protocols were approved by the Ethics Committees of the Universities of Exeter and Pretoria and complied with regulations stipulated in The Association for the Study of Animal Behaviour (ASAB) Guidelines for Use of Animals in Research.

### Monitoring provisioning behavior

To identify individuals during the recording of nestling provisioning events, group members were captured from their roost chambers (details below) during the incubation period and marked on the vent with a unique dye-mark. The dominant female was left unmarked to minimize disturbance during incubation, but could still be distinguished from other group members by being the only unmarked bird within her group (only resident group members provision offspring). To record provisioning events, a Panasonic SDR-S50 camcorder attached to a tripod (approximately 0.5 meters in height) was placed on the ground beneath the entrance to the active nest two days before recording commenced (to allow the birds to habituate to it). On the days of provisioning monitoring, the recordings (approximately 3 hours in the duration) were started between 06:15 and 07:54, with this start time being adjusted through the season to maintain an approximately constant time offset from sunrise. Provisioning videos were collected in this way for all focal breeding attempts (n = 37 broods across 30 social groups) on two mornings between Days 6 and 9 inclusive of the nestling provisioning period (typically for the two consecutive mornings of Days 7 and 8; nestlings fledged from day 20). This approach yielded a mean total duration of provisioning video of 6.08 hours (range 4.05 – 8.12 hours) per breeding attempt. Video recordings were transcribed using VLC Media Player version 2.2, with the observer recording, for each provisioning visit, the identity of the bird visiting the nest (determined via their distinct dye mark and bill color, which reveals their sex) and the duration of time that they spent within the nest (the time elapsed between passing in and out of the enclosed nest structure; hereafter ‘Provisioning visit duration’). Prior work on this study population using within-nest cameras has shown that all nest visits during the nestling age window studied here entail the delivery of a single prey item to the brood, unless the visiting bird is carrying a feather or grass in which case no food is delivered (Walker, 2016). We therefore excluded such feather- or grass-carrying nest visits from our provisioning visit records. From the transcribed data for each focal brood we then calculated two provisioning trait values for each adult group member: (i) ‘Provisioning rate’ (feeds / hr) was calculated as the total number of provisioning visits that the bird conducted over the two monitored mornings divided by the total duration of video collected over those two mornings, and (ii) ‘Mean Visit Duration’ (minutes) was calculated as the mean duration of all provisioning visits conducted by the focal bird over the two monitored mornings.

### Bird capture and blood sampling

To obtain a matched blood sample for prolactin measurement, we attempted to capture and blood sample all adult (> 1 year old at the time of sampling) birds within the monitored brood’s social group on the evening of the second day of provisioning behavior recording. Birds were captured individually at night from the woven roost chambers within their group’s territory (Cram et al., 2015a) by flushing individuals into a custom-made capture bag. All captures, dye-marking and blood sampling were conducted by a single investigator (LW). Birds were then immediately returned to a roost chamber within their territory to pass the remainder of the night.

Upon capture, a blood sample (*c*. 140 µL) was taken from the brachial vein of the bird using a 26g needle and heparinized capillary tubes. Captures occurred soon after dusk, once the birds were roosting in their woven chambers. Time of capture was recorded (to allow us to fit the time lag from sunset to capture as a covariate predictor in our prolactin analyses, in case of diel variation and/or effects of the time elapsed since roosting on a bird’s prolactin levels), along with the time lag between capture and the completion of blood sampling (mean ± standard deviation [S.D.] = 3.12 ± 0.73 minutes [range 1.65 – 4.78 minutes]; to allow us to control for potential effects of capture stress on prolactin levels in our statistical models). Blood samples were immediately centrifuged in the field (12,000 g for 3 minutes; Haematospin 1400; Hawksley Medical and Laboratory Equipment, Lancing, UK) and the plasma was drawn off and stored in a cryovial on ice until it could be transferred to liquid nitrogen on return from the field (mean ± S.D. time lag from sample collection to storage on liquid nitrogen = 148 ± 63 min). At the end of the field season, samples were transferred to the UK on dry ice and then stored at −80 degrees Celsius until analysis for prolactin.

### Prolactin Radioimmunoassay

The prolactin assay was carried out in July 2014 at the Roslin Institute (University of Edinburgh, Easter Bush, Midlothian, Scotland, UK). Plasma prolactin levels were measured using a highly specific heterologous micro-radioimmunoassay of donkey anti-rabbit serum to European starling (*Sturnus vulgaris*) prolactin (Sharp antibody code 44/2). Prolactin was radiolabeled with iodine^125^ using chloramine-T. 168 (out of a total of 208) samples were assayed in duplicate, and the remaining 40 samples were assayed as singletons (not all samples assayed were for use in this study). All samples were measured in a single assay, in which the intra-assay coefficient of variation for the duplicate samples was 3.31%.

### Statistical methods

The above methods yielded a final data set of matched provisioning trait data (estimated from the focal bird’s average performance over two mornings of provisioning recordings; see above) and circulating prolactin levels (when sampled on the evening of the second day of provisioning monitoring) for 70 different adult birds, each sampled once (for all traits), while feeding a total of 37 broods across 30 social groups. While some analyses utilized the whole data set (i.e. n = 70 adults birds each sampled once for all traits), others used subsets of it (e.g. when focusing only on dominants engaged in parental care or subordinates engaged in cooperative helping behavior), and so the sample sizes for each analysis are reported within the relevant results section and model output table. As mean provisioning visit duration data were only available for birds that had a non-zero provisioning rate, the sample sizes for mean visit duration analyses were sometimes smaller than those for provisioning rate (see results).

All statistical models and visualizations were carried out in R (version 4.1.0; R Core Team). Mixed effects modelling was conducted using the R package ‘lme4’ (Bates et al., 2015) We conducted our statistical modelling using a full model approach, in which we (i) specified a full model containing both the primary fixed effect predictors of interest and covariates whose potential effects we also wished to control for, and then (ii) tested the effects of these predictors in that context, without any model selection or simplification. The statistical significance of a given predictor was assessed by using a likelihood ratio test to determine the significance of the change in the explanatory power of the full model when the focal predictor was dropped from the full model. This conservative approach ensures that the significance of all predictors is assessed while controlling for the potential effects of the other predictors specified in the full model, regardless of whether those other predictors themselves have significant effects. The specific modelling exercises conducted for each results section are described below.

#### 1. Are differences in the provisioning behavior of dominant females and males (engaged in parental care) mirrored by differences in their circulating prolactin levels?

We used two separate mixed effects models with Gaussian error structure to model the causes of variation in (i) the provisioning rates and (ii) the mean provisioning visit durations of dominant birds (i.e. engaged in parental care). The two modeling exercises began with an identical full model structure. In addition to the primary predictor of interest, ‘parent class’ (dominant female or dominant male), we fitted the following terms as fixed effect predictors: brood size (the brood size being fed), adult group size (the number of adult group members during the focal nestling provisioning period) and year (a two-level factor capturing the year in which sampling occurred; 2013 or 2014). We fitted both social group ID and brood ID (the identity of the brood being fed) as random effects, retaining them in the model structure regardless of the degree of variance that they explained.

We then used a third mixed effects model with Gaussian error structure to model the causes of variation in the circulating prolactin levels of these same dominant birds, starting with a full model structure containing the same fixed and random effect predictors as the provisioning trait models just described, but with the addition of two further fixed effects to account for potential methodological effects on prolactin concentrations: (i) the time lag from sunset to the bird’s capture for blood sampling (to allow for the possibility of diel variation in prolactin levels and/or changes in prolactin levels once the birds entered their roosts) and (ii) the time lag from capture to the completion of blood sampling (to allow for possible effects of capture stress on circulating prolactin levels).

#### 2. Are differences in the provisioning behavior of natal and immigrant subordinates (engaged in alloparental helping behavior) mirrored by differences in their circulating prolactin levels?

We then used two mixed effects models with Gaussian error structure to model the causes of variation in the provisioning rates and circulating prolactin levels of subordinate birds engaged in alloparental helping behavior. We fitted the same set of fixed and random effect predictors to these models as were fitted to the corresponding provisioning rate and prolactin level models conducted for dominant birds (see above) with two exceptions: (i) in place of ‘parent class’ we fitted ‘helper class’, reflecting whether the focal bird was a natal subordinate or an immigrant subordinate, and (ii) here only brood ID was fitted as a random effect (social group ID was not, as all subordinate birds from any given social group were sampled while feeding the same single brood, leaving brood ID and social group ID with identical structure). We did not model the causes of variation in the mean provisioning visit durations of subordinates as too few immigrant subordinates actually provisioned the focal broods, leaving us with an insufficient sample size of measures of the provisioning visit durations of this bird class.

#### 3. Does continuous variation in prolactin levels predict variation in provisioning behaviour?

To investigate whether natural variation in prolactin levels predicted continuous variation in the birds’ nestling provisioning rates and mean provisioning visit durations at the population level (i.e. when all bird classes were combined) we conducted two mixed effect models (one for each provisioning trait response term), with circulating prolactin concentration as the sole fixed effect predictor (as we have not hypothesized specific mechanisms by which other variables might impact provisioning traits independent of prolactin levels) and social group ID and brood ID as random effects. Mean provisioning visit duration was logarithm transformed for analysis, to normalize model residuals.

Inspection of the patterns of the mean prolactin levels and provisioning trait values of the four different bird classes (i.e. dominant females, dominant males, natal subordinates and immigrant subordinates) suggested that any such continuous relationship between prolactin levels and provisioning trait values at the population level across all bird classes could be driven principally by the variation in these traits *among* the bird classes (Figures 3a & 3c). In order to then investigate whether the more limited variation in circulating prolactin levels *within* bird classes predicted the *within*-class variation in provisioning trait values, we first mean-centered each birds’ prolactin level and provisioning trait values (the log transformed values in the case of mean provisioning visit duration) around the mean value of the focal trait for birds of their class (by subtracting from it the mean value of the focal trait for their bird class). We then conducted two mixed effects models (one for each mean-centered provisioning trait response term), with mean-centered circulating prolactin concentration as the sole fixed effect predictor and social group ID and clutch ID as random effects.

## RESULTS

### 1. Are differences in the provisioning behavior of dominant females and males (engaged in parental care) mirrored by differences in their circulating prolactin levels?

Analyzing the provisioning behavior of dominant birds engaged in parental care (n = 46 dominants, 20 females and 26 males, feeding 31 broods at 28 social groups) revealed that dominant females feed offspring at significantly higher rates than dominant males (Figure 1a; parent class effect, for dominant male relative to dominant female, ± S.E. = +5.11 ± 0.57; P < 0.001; Table 1) and also show significantly longer mean provisioning visit durations (Figure 1b; parent class effect, for dominant male relative to dominant female, ± S.E. = +1.29 ± 0.31; P < 0.001; Table 2). There was also evidence that dominant birds fed larger broods at significantly higher rates (brood size effect ± S.E. = +1.08 ± 0.50; P = 0.035; Table 1). There was no compelling evidence that the provisioning rates or mean visit durations of dominant birds were associated with either the year of study or adult group size (Tables 1 & 2).

**Figure 1.**
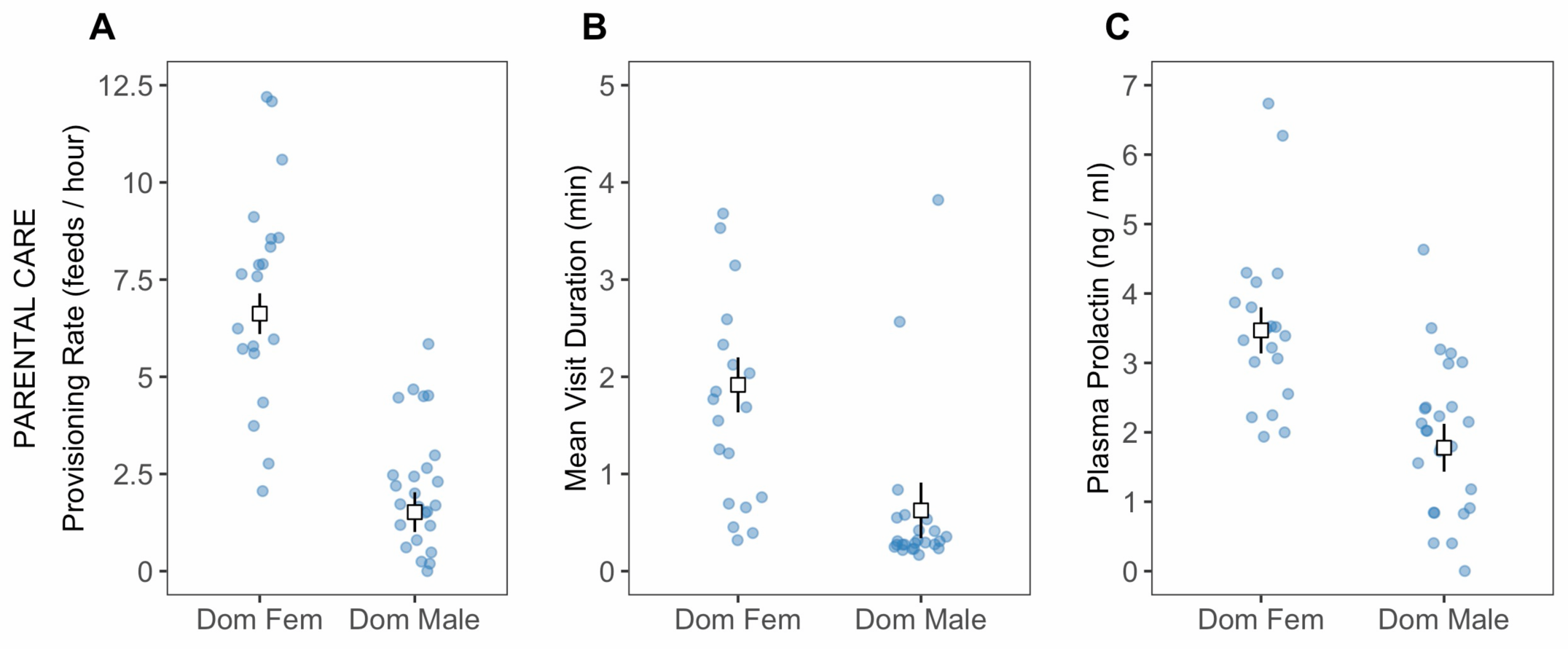
The (a) provisioning rates, (b) mean provisioning visit durations, and (c) circulating prolactin concentrations of dominant females (Dom Fem) and dominant males (Dom Male) engaged in parental care. Squares present the predicted means (± S.E.) from the full model for the relevant trait (Tables 1, 2 and 3 respectively), while controlling for the effects of all other variables in the full model. The predicted means were calculated with all continuous predictors set to their mean values and for the 2013 category of the Season factor. The points show the raw data points.

**Table 1:** Modelling the effects of Parental Sex on Nestling Provisioning Rate during Parental care. N = 46 dominant birds, 20 female and 26 male, feeding 31 broods at 28 social groups. The ‘Parent class’ effect size is for the dominant female relative to the dominant male. The ‘Year’ effect size is for the 2014 field season relative to the 2013 field season. Significant effect sizes (on the basis of likelihood ratio tests when comparing the full model to the full model without the focal term) are highlighted in bold. SE = Standard Error; LRT = Likelihood ratio test and associated P value.

|  | Estimate | SE | t | LRT | P |
| --- | --- | --- | --- | --- | --- |
| (Intercept) | -0.461 | 1.252 | -0.368 |  |  |
| **Parent class**<br>(female > male) | **5.106** | **0.568** | **8.984** | **46.611** | **<0.001** |
| Year | 1.027 | 0.638 | 1.608 | 2.516 | 0.113 |
| Group size | 0.050 | 0.276 | 0.181 | 0.033 | 0.856 |
| **Brood size** | **1.084** | **0.501** | **2.164** | **4.461** | **0.035** |

**Table 2:** Modelling the effects of Parental Sex on Mean Nestling Provisioning Visit Duration during Parental care. N = 44 dominant birds, 20 female and 24 male, feeding 31 broods at 28 social groups; the sample size for this analysis was slightly lower than that for the Provisioning Rate analysis (Table 1) as 2 dominant males did not provision their brood, leaving us without a measure of their mean visit duration. The ‘Parent class’ effect size is for the dominant female relative to the dominant male. The ‘Year’ effect size is for the 2014 field season relative to the 2013 field season. Significant effect sizes (on the basis of likelihood ratio tests when comparing the full model to the full model without the focal term) are highlighted in bold. SE = Standard Error; LRT = Likelihood ratio test and associated P value.

|  | Estimate | SE | t | LRT | P |
| --- | --- | --- | --- | --- | --- |
| Intercept | 0.960 | 0.720 | 1.333 |  |  |
| **Parent class**<br>(female > male) | **1.292** | **0.314** | **4.119** | **14.351** | **<0.001** |
| Year | -0.091 | 0.348 | -0.262 | 0.069 | 0.793 |
| Group size | 0.032 | 0.153 | 0.210 | 0.044 | 0.834 |
| Brood size | -0.253 | 0.285 | -0.887 | 0.780 | 0.377 |

**Table 3:** Modelling the effects of Parental Sex on Circulating Prolactin level during Parental care. N = 46 dominant birds, 20 female and 26 male, feeding 31 broods at 28 social groups (identical to the nestling provisioning rate analysis in Table 1, as all birds were sampled for both traits in the same contexts). The ‘Parent class’ effect size is for the dominant female relative to the dominant male. The ‘Year’ effect size is for the 2014 field season relative to the 2013 field season. Significant effect sizes (on the basis of likelihood ratio tests when comparing the full model to the full model without the focal term) are highlighted in bold. SE = Standard Error; LRT = Likelihood ratio test and associated P value. Sunset to capture lag = time elapsed between sunset and capture (to account for potential circadian variation in prolactin levels). Capture to bleed lag = time elapsed between first contact with the roost chamber and blood sample (to allow for the possibility of a prolactin stress response).

|  | Estimate | SE | t | LRT | P |
| --- | --- | --- | --- | --- | --- |
| (Intercept) | 0.731 | 1.199 | 0.610 |  |  |
| **Parent class**<br>(female > male) | **1.691** | **0.261** | **6.487** | **23.196** | **<0.001** |
| Year | 0.328 | 0.446 | 0.735 | 0.378 | 0.539 |
| Group size | 0.271 | 0.180 | 1.509 | 2.217 | 0.137 |
| Brood size | 0.298 | 0.320 | 0.931 | 0.783 | 0.376 |
| Sunset to capture lag | 0.000 | 0.003 | -0.002 | 0.000 | 0.998 |
| Capture to bleed lag | -0.001 | 0.004 | -0.295 | 0.087 | 0.768 |

Analyzing the circulating prolactin levels of dominant birds during the provisioning periods analyzed above (again, n = 46 dominants, 20 females and 26 males, feeding 31 broods at 28 social groups) revealed evidence that dominant females also have significantly higher circulating prolactin levels than dominant males (Figure 1c; parent class effect, for dominant male relative to dominant female, ± S.E. = +1.69 ± 0.26; P < 0.001; Table 3). There was no compelling evidence that the prolactin levels of dominants were associated with group size, brood size, the time lag from sunset to capture or the time lag from capture to blood sampling (Table 3). Note that our full model approach ensured that any effects of these latter predictors were controlled (regardless of their significance) when assessing the effects of other terms.

### 2. Are differences in the provisioning behavior of natal and immigrant subordinates (engaged in alloparental helping behavior) mirrored by differences in their circulating prolactin levels?

Analyzing the provisioning behavior of subordinate birds engaged in cooperative helping behavior (n = 24 subordinates, 17 natal and 7 immigrant, feeding 16 broods at 16 social groups) revealed evidence that subordinates within their natal groups feed offspring at higher rates than immigrant subordinates (Figure 2a; helper class effect, for natal subordinates relative to immigrant subordinates, ± S.E. = +1.61 ± 0.33; P < 0.001; Table 4). There was also evidence that helpers fed larger broods at significantly higher rates (brood size effect ± S.E. = +1.06 ± 0.35; P = 0.007; Table 4). There was no compelling evidence that helper provisioning rates were associated with either group size or the year of study (Table 4). No analysis of the provisioning visit durations of subordinates was conducted as an insufficient number of subordinate immigrants ever provisioned the broods (see Figure 2a).

**Figure 2.**
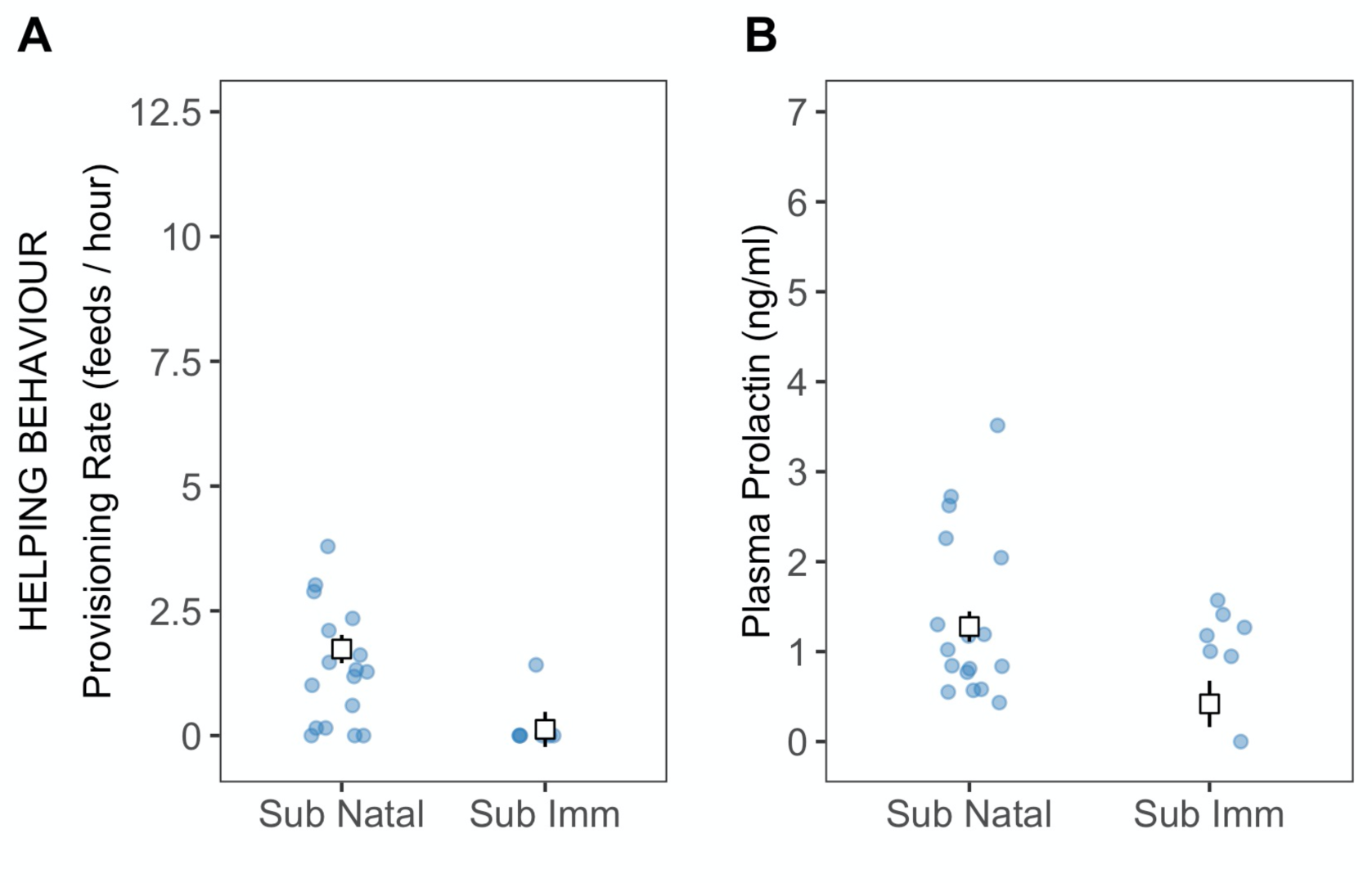
The (a) provisioning rates, and (b) circulating prolactin levels of natal subordinates (Sub Natal) and immigrant subordinates (Sub Imm) engaged in cooperative helping behavior, feeding the broods of the dominant male and female. Y axis scales match those in Figure 1 to facilitate comparison. Squares present the predicted means (± S.E.) from the full model for the relevant trait (Tables 4 and 5 respectively), while controlling for the effects of all other variables in the full model. The predicted means were calculated with all continuous predictors set to their mean values and for the 2013 category of the Season factor. The points show the raw data points. No analysis of provisioning visit durations was conducted as an insufficient number of subordinate immigrants ever provisioned the broods (see panel 2a).

**Table 4:** Modelling the effects of Helper Class on Nestling Provisioning Rate during Alloparental Helping Behaviour. N = 24 subordinate birds, 17 natal and 7 immigrant, feeding 16 broods at 16 social groups. The ‘Helper class’ effect size is for natal subordinates (those residing within their natal groups) relative to immigrant subordinates (those that have dispersed to another group). The ‘year’ effect size is for the 2014 field season relative to the 2013 field season. Significant effect sizes (on the basis of likelihood ratio tests when comparing the full model to the full model without the focal term) are highlighted in bold. SE = Standard Error; LRT = Likelihood ratio test and associated P value.

|  | **Estimate** | **SE** | **t** | **LRT** | **P** |
| --- | --- | --- | --- | --- | --- |
| Intercept | -0.297 | 1.097 | -0.271 |  |  |
| **Helper class**<br>(natal > immigrant) | **1.610** | **0.334** | **4.826** | **15.858** | **<0.001** |
| Year | -0.810 | 0.435 | -1.861 | 3.173 | 0.075 |
| Group size | -0.274 | 0.226 | -1.212 | 1.415 | 0.234 |
| **Brood size** | **1.057** | **0.354** | **2.986** | **7.166** | **0.007** |

**Table 5:** Modelling the effects of Helper Class on Circulating Prolactin level during Alloparental Helping Behaviour. . N = 24 subordinate birds, 17 natal and 7 immigrant, feeding 16 broods at 16 social groups (identical to the nestling provisioning rate analysis in Table 4, as all birds were sampled for both traits in the same contexts). The ‘Helper class’ effect size is for natal subordinates (those residing within their natal groups) relative to immigrant subordinates (those that have dispersed to another group). The ‘year’ effect size is for the 2014 field season relative to the 2013 field season. Significant effect sizes (on the basis of likelihood ratio tests when comparing the full model to the full model without the focal term) are highlighted in bold. SE = Standard Error; LRT = Likelihood ratio test and associated P value. Sunset to capture lag = time elapsed between sunset and capture (to account for potential circadian variation in prolactin levels). Capture to bleed lag = time elapsed between first contact with the roost chamber and blood sample (to allow for the possibility of a prolactin stress response).

|  | **Estimate** | **SE** | **t** | **LRT** | **P** |
| --- | --- | --- | --- | --- | --- |
| Intercept | 1.255 | 0.910 | 1.378 |  |  |
| **Helper class**<br>(natal > immigrant) | **0.860** | **0.281** | **3.054** | **7.881** | **0.005** |
| **Year**<br>(2014 > 2013) | **0.848** | **0.370** | **2.294** | **4.757** | **0.029** |
| **Group size** | **-0.365** | **0.126** | **-2.901** | **7.152** | **0.007** |
| Brood size | -0.175 | 0.227 | -0.770 | 0.586 | 0.444 |
| Sunset to capture lag | 0.002 | 0.003 | 0.709 | 0.498 | 0.480 |
| Capture to bleed lag | 0.004 | 0.003 | 1.561 | 2.320 | 0.128 |

Analyzing the circulating prolactin levels of subordinate birds during the provisioning periods analyzed above (again, n = 24 subordinates, 17 natal and 7 immigrant, feeding 16 broods at 16 social groups) revealed that natal subordinates also have significantly higher prolactin levels than immigrant subordinates (Figure 2b; helper class effect, for natal subordinates relative to immigrant subordinates, ± S.E. = +0.86 ± 0.28; P = 0.005; Table 5). There was also evidence that subordinate prolactin levels were significantly higher in the second year of study (year effect, for 2014 relative to 2013, ± S.E. = +0.85 ± 0.37; P = 0.029; Table 5) and in smaller groups (Group size effect ± S.E. = −0.37 ± 0.13; P = 0.007; Table 5). There was no compelling evidence that subordinate prolactin levels were associated with brood size or the time lags from sunset to capture and from capture to blood sampling (Table 5). Note that our full model approach ensured that any effects of these latter predictors were controlled (regardless of their significance) when assessing the effects of other terms.

### 3. Does continuous variation in prolactin levels predict variation in provisioning behaviour?

Our analysis at the population level, including birds of all classes, revealed that a bird’s circulating prolactin level strongly and significantly positively predicts both (i) its provisioning rate (Figure 3a; effect size ± S.E. = 1.20 ± 0.23; P < 0.001; n = 70 birds feeding 37 broods at 30 social groups) and (ii) its mean provisioning visit duration (Figure 3c; effect size ± S.E. = 0.32 ± 0.076; P < 0.001; n = 59 birds feeding 36 broods at 30 social groups). Plotting out the mean prolactin levels and provisioning trait values of the different bird classes (Figure 3a & 3c), reveals that both of these population-level relationships between prolactin and provisioning traits are driven in large part by the among-bird-class differences in prolactin levels being mirrored by parallel among-bird-class differences in mean provisioning rate (Figure 3a) and mean provisioning visit duration (Figure 3c). Indeed, after mean-centering each bird’s prolactin level and provisioning trait values around the focal trait’s mean value for their bird class, we found no evidence that within-bird-class variation in prolactin levels predicted within-bird-class variation in either provisioning rate (Figure 3b; effect size ± S.E. = −0.15 ± 0.21; P = 0.50) or mean provisioning visit duration (Figure 3d; effect size ± S.E. = 0.049 ± 0.081; P = 0.55).

**Figure 3.**
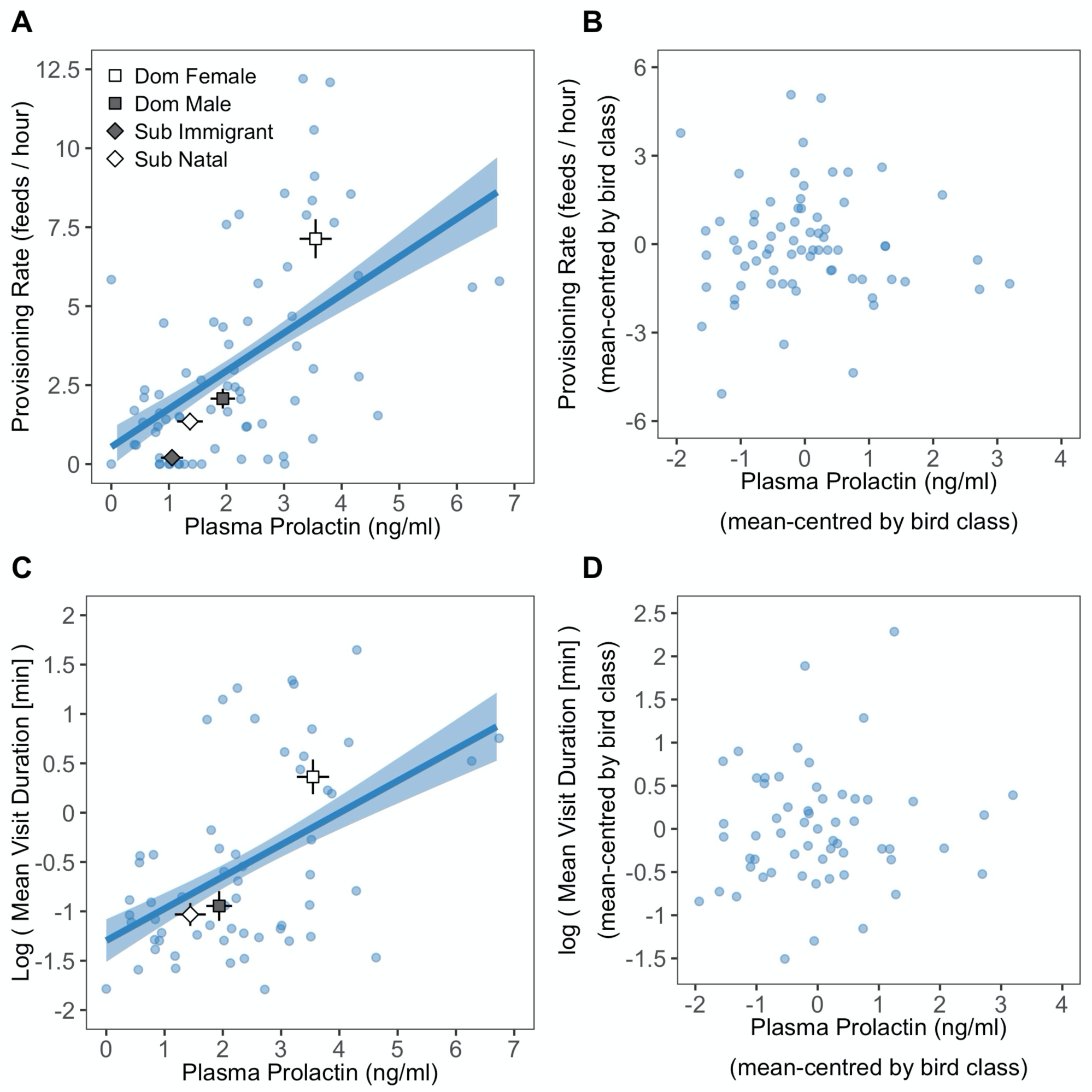
At the population level, considering all bird classes together, natural variation in circulating prolactin levels predicts variation in both (**a**) provisioning rate and (**c**) mean provisioning visit duration. These relationships are driven principally by differences among the mean trait values of the different focal bird classes (presented ± S.E. by the squares and diamonds within panels **a** and **c**; see legend within panel **a**). Follow-up analyses revealed no evidence that variation in prolactin levels *within* these bird classes predicted within-bird-class variation in either (**b**) provisioning rate or (**d**) mean provisioning visit duration. In panels **a** and **c** the line and shaded ribbon present the predicted mean relationship and its standard error, while the shaded circular points within all panels present the raw data points.

## DISCUSSION

This study investigated the hypothesis that prolactin plays a role in the regulation of nestling provisioning, both among dominant birds (engaged in parental care) and non-breeding subordinate birds (engaged in cooperative helping behavior), in cooperatively breeding white-browed sparrow weaver societies. Among dominants engaged in parental care, we found that the dominant female (the mother) fed offspring at higher rates, made longer provisioning visits and had higher circulating prolactin levels than the dominant male (typically the father). Among subordinates, we found that natal subordinates helped to feed offspring at higher rates and had higher circulating prolactin levels than immigrant subordinates. Indeed, when all bird classes were combined, we found that continuous variation in the circulating prolactin levels of the birds predicted continuous variation in their provisioning rates and mean provisioning visit durations. These patterns appear to be driven principally by correlated differences among the four different bird classes in their prolactin levels and provisioning traits. We found no evidence that the more limited variation in circulating prolactin levels within the different bird classes predicted the more limited within-class variation in their provisioning traits. Together, these findings are broadly consistent with the hypothesis that parental care and cooperative helping behavior are regulated by a common underlying mechanism and that prolactin plays a role in that pathway, and highlight the need for experimental studies to now probe the causality and nature of any role for prolactin. Below, we discuss potential explanations for these findings, the different roles that prolactin could conceivably play in the regulation of parenting and cooperative helping in this species, and the wider implications of our findings for mechanistic and evolutionary research on cooperative behavior.

While our findings are consistent with the hypothesis that parental care and helping behavior are regulated by a common mechanism in which prolactin plays a role, the lack of a relationship between within-class variation in prolactin levels and provisioning traits, coupled with the correlative nature of our findings, leave it important to consider the range of possible roles that prolactin could play in the regulation of provisioning behavior in this species. At least three main possibilities exist, which will require careful experimentation to tease apart. First, it is possible that circulating prolactin is one key regulator of continuous variation in individual contributions to offspring provisioning, both among parents and helpers. While most of our findings are consistent with this hypothesis, the absence of evident relationships between within-class variation in prolactin levels and provisioning traits complicates this view. However, the lack of evident *within*-class relationships could be attributable simply to a major source of variation in both traits (*among*-class variation) having been factored out at this stage of the analysis, leaving these within-class analyses seeking relationships between the more limited *within*-class variation in both traits, which could be readily obscured by a number of mechanisms. First, difficulties with the synchronous and accurate assessment of both prolactin levels and provisioning rates could have yielded noise in the data set that precluded the detection of these more subtle prolactin-provisioning relationships. While we sampled birds for prolactin on the evening following the morning provisioning-monitoring session (a time lag comparable to, or shorter than, those of similar studies; e.g. Duckworth et al., 2003; Ouyang et al., 2011), individuals may have differed in the way that their prolactin levels changed during the day, leaving their evening prolactin levels only a modest proxy for those while provisioning. The focal birds also varied in the timing of blood sampling, and while our analyses did not detect any overall effects on prolactin levels of the time lag from capture to sampling, any individual variation in the prolactin stress response (if this species shows one; Krause et al., 2015) could have further decoupled the assessed prolactin levels from those during provisioning. Second, even if prolactin levels were a key regulator of continuous variation in provisioning rates, alternative mechanisms are also expected to impact provisioning rates potentially independent of circulating prolactin levels, leaving the relationship between natural variation in prolactin levels and provisioning behavior potentially weak in the first place (Schoech et al., 1998; Angelier et al., 2016). Key among these could be (i) variation in other components of a prolactin-mediated pathway (such as inter-individual and temporal variation in the density of prolactin receptors; Zhou et al., 1996; Ohkubo et al., 1998; Angelier et al., 2016), as well as (ii) mechanisms that may impact provisioning behavior via prolactin-independent pathways (e.g. the effects of circulating testosterone; Schoech et al., 1998; Angelier et al., 2016). Third, even if prolactin levels alone determined provisioning ‘motivation’, the extent to which variation in provisioning motivation was reflected in provisioning rates would depend upon the prey capture skills of the focal bird and the environmental availability of prey. Indeed, all points considered, it is arguably mechanistically naïve to expect particularly fine-grained associations between the levels of a single hormone and behavior to be evident in natural populations even where a causal link exists between the two. To now robustly test the hypothesis that prolactin regulates continuous variation in the magnitude of both parental and helper contributions to offspring provisioning, there is a need to experimentally elevate the circulating prolactin levels of actively provisioning birds whose natural prolactin levels are not at the upper end of the physiological range (dominant males and natal subordinates may serve this purpose well; Figure 3a). This manipulation would allow one to test the key prediction that an increase in the prolactin levels of an actively provisioning bird will cause it to increase its provisioning rate; a prediction that to our knowledge has yet to be tested in either a parenting or helping context (the few experimental elevations of endogenous prolactin secretion in a provisioning context to date have focussed on the establishment of provisioning in non-provisioning birds rather than its quantitative variation within actively provisioning birds; e.g. Badyaev and Duckworth, 2005).

A second potential explanation for the balance of our findings is that prolactin could instead play a causal role in the onset and maintenance of provisioning behavior among parents and helpers, without playing a role in the quantitative regulation of contributions to provisioning among actively provisioning birds (Angelier et al., 2016). For example, a threshold level of prolactin may be required for the onset and/or maintenance of provisioning behavior (Angelier et al., 2006; Boos et al., 2007). Under this scenario, the higher prolactin levels of natal subordinates and dominant birds, relative to immigrant subordinates, could be causally responsible for the former bird classes engaging in provisioning while the latter typically does not. This could be the case without prolactin playing any causal role in regulating continuous variation in the provisioning rates of actively provisioning birds; a scenario that could account for the lack of within-class correlations between prolactin levels and provisioning behavior. The elevated prolactin levels of dominant females (relative to dominant males and natal subordinates) could conceivably be a downstream consequence of either a role for prolactin in incubation (Buntin, 1996; Sharp et al., 1998; Khan et al., 2001; as dominant females are the sole incubator in this species) and/or their differential exposure to offspring cues during the nestling period (which can increase prolactin secretion; Hall, 1987; Sharp et al., 1998), given their markedly higher provisioning rates and mean visit durations than other classes. The hypothesis that prolactin maintains provisioning behavior but does not quantitatively regulate contributions to it could now be tested by (i) experimentally elevating the prolactin levels of subordinate immigrants, to test the prediction that this would cause these typically non-provisioning birds to commence provisioning behavior (e.g. see Badyaev and Duckworth (2005) for a demonstration of this transition in the context of parental nestling feeding), (ii) experimentally reducing the prolactin levels of the actively-provisioning classes to test whether this eliminates provisioning behavior (e.g. Smiley and Adkins-Regan, 2018), and (iii) experimentally elevating the prolactin levels of actively provisioning dominant males and/or natal subordinates (the manipulation proposed in the previous paragraph), as doing so should *not* increase their provisioning rates if prolactin merely maintains provisioning behavior without regulating contributions to it.

Given the correlative nature of our findings, it is also conceivable that prolactin plays no causal role in the onset, maintenance or quantitative regulation of parenting and/or cooperative helping in this species (despite experimental evidence of causal effects on parenting in other species; see Introduction; Angelier et al., 2016). In this scenario, one might attribute the evident associations between prolactin and provisioning to a ‘reverse causal’ relationship, in which provisioning interactions with offspring stimulate prolactin release (Hall, 1987; Sharp et al., 1998). However, such a reverse causal argument alone cannot readily account for our findings in their entirety, as within-class variation in provisioning rates and mean provisioning visit durations were not evidently associated with prolactin levels (though, again, the lack of such an association could be attributable to challenges with accurately and simultaneously quantifying hormone and behavior; see above). When considering whether our findings could be attributable solely to effects of provisioning on prolactin levels (i.e. in the absence of any effect of prolactin on provisioning), it is worth considering why selection would have left prolactin levels sensitive to offspring interactions in the first place. Arguably the most plausible explanation is that this mechanism plays a role in a feedback loop in which a causal relationship exists in both directions: if prolactin did establish, maintain and/or regulate care, selection may have favored regulating prolactin secretion according to offspring interactions in order to maintain care as long as offspring survive and/or regulate care according to offspring viability or need (Hall, 1987; Sharp et al., 1998; Angelier et al., 2016). As such, where offspring cues do stimulate prolactin release, such a relationship might generally be expected to occur alongside causal effects of prolactin on care. While the experiments outlined above would shed light on the causality of the prolactin-provisioning associations detected here, wider investigations are also needed to probe the role, if any, that such a feedback loop (with causal relationships in both directions) may play in the maintenance and/or regulation of cooperative care.

While experimental tests of causality are needed, our findings are broadly consistent with the hypothesis that pre-existing mechanisms that regulated parental care in ancestral bi-parental species were co-opted for the regulation of cooperative helping behavior on the evolution of cooperative breeding. The often-overlooked possibility that parenting and cooperative helping are indeed regulated by a common mechanism has important evolutionary implications. Explanations for the evolution, maintenance and optimization of cooperative behavior typically focus on the roles of the fitness benefits and costs of cooperation *per se* (Hamilton, 1964; Cockburn, 1998; West et al., 2007; Capilla-Lasheras et al., 2021). However, if cooperation and parenting are regulated by a common underlying mechanism, it is conceivable that this shared regulatory architecture for care giving is shaped as much by the payoffs from its outcomes in a parental context as by the payoffs from its outcomes in a cooperative helping context. While selection might independently optimize parental and non-breeding helper caring strategies (e.g. via the evolution of an entirely context-dependent caring strategy), it is conceivable that mechanistic constraints preclude their independent optimization. For example, genetic variants that modified sensitivity to begging could conceivably impact the expression of both parental care and cooperative helping, yielding scope for intra-locus genetic conflict to constrain the independent optimization of both parental care and cooperative helping (Pennell et al., 2018; see also the conceptual parallels with sexual conflict: Stewart et al., 2010; Pennell and Morrow, 2013). Where this is the case, attempts to understand the evolutionary origins, maintenance and optimization of cooperative behavior may require attention to the extent to which genetic correlations exist between parental and cooperative behavior. Notably, our findings suggest that cooperative helping behavior in sparrow-weaver societies is not maintained by selection *solely* because a genetic correlation with parenting has precluded the evolution of ‘non-helping’ (see Brown and Vleck, 1998 for a similar debate), because a context-dependent helping strategy does appear to have evolved. Subordinates routinely help while within their natal groups (where they are closely related to the broods that they help to rear; Harrison et al., 2013a), but typically cease to do so following immigration into another group (where they are typically unrelated to broods, reducing the potential indirect fitness payoff from helping; Harrison et al., 2013a). While endocrine research on cooperative breeders has historically focused principally on the proximate causes of the rank-related reproductive disparities that typify such societies (Schoech et al., 2004; Young et al., 2006; 2008), a renewed focus on the endocrinology of care in cooperative breeders (Schoech et al., 2004; Soares et al., 2010; Dantzer et al., 2017; 2019) would now help to shed light on the extent to which parenting and cooperation are indeed regulated by shared underlying pathways.

## CONCLUSION

Our findings lend new support to the hypotheses that helping behavior in cooperatively breeding societies has shared mechanistic underpinnings with parental care, and that prolactin plays a key role in this pathway (see also Vleck et al., 1991; Schoech et al., 1996; Khan et al., 2001; Carlson et al., 2006). Our findings and their complexity highlight the need for experimental studies to investigate both the causality and nature of the relationship between prolactin and provisioning in this species, in both parental and cooperative helping contexts. Our findings also highlight that attempts to understand the evolution of cooperative helping may benefit from attention to the possibility of constraints on the independent optimization of cooperative helping and parenting. Our study has implications too for the growing interest in the mechanistic origins of consistent individual differences in cooperative helping behavior (Sanderson et al., 2015; Dantzer et al., 2019). Specifically, our findings highlight that such differences could arise from consistent individual differences within the pathway by which prolactin acts (e.g. via differences in prolactin secretion and/or reception; Ohkubo et al., 1998; Zhou et al., 1996).

## ACKNOWLEDGMENTS

We dedicate this manuscript to our co-author Peter Sharp (P.J.S.), whose openness and interest in this project were key to its inception, and whose wider contributions to avian endocrinology we celebrate. Any errors are our own, as he did not have a chance to comment on the final manuscript. We are grateful to D. O’Connell for support with data collection, D. L. Cram for assistance in training in bird capture and blood sampling, N.C. Bennett for invaluable logistical support, J.L. McDonald and K.J. Walker for comments on an earlier draft, and P. Capilla-Lasheras for statistical advice. We also thank E.O. and Son and all at Tswalu Kalahari Reserve for access to our field site and exceptional logistical support (with particular gratitude to Cornelia le Roux and Dylan Smith), and the Northern Cape Conservation Authority for granting permission to conduct our research. L.A.W. was funded by a NERC PhD studentship (NE/J500185/1). J.E.Y. was funded by the BBSRC (BB/H022716/1). S.L.M. was funded by Roslin Institute Strategic Grant funding from the BBSRC (BB/P013759/1). A.J.Y. was funded by BBSRC David Phillips and NERC Blue Skies Research Fellowships (BB/H022716/1 and NE/E013481/1).

